# Pairwise genomic alterations identify prognostic tumor states in multiple cancer types

**DOI:** 10.64898/2026.04.14.718563

**Authors:** Masroor Bayati, Jigyansa Mishra, Laura Luhari, Maria Farina-Morillas, Jüri Reimand

**Affiliations:** Computational Biology Program, Ontario Institute for Cancer Research, Toronto, ON, Canada; Department of Medical Biophysics, University of Toronto, Toronto, ON, Canada; Department of Molecular Genetics, University of Toronto, Toronto, ON, Canada; Department of Chemistry and Biotechnology, Tallinn University of Technology, Tallinn, Estonia; Cancer Computational Biology Group, Vall d’Hebron Institute of Oncology (VHIO), Barcelona, Spain

## Abstract

Genomic models of cancer prognosis usually rely on individual genomic alterations, potentially overlooking clinically meaningful combinations of events. We analyzed genomic and clinical data from nearly 10,000 primary tumors across major cancer types to identify prognostic genomic interactions (PGIs), defined as pairs of genomic alterations whose joint status was associated with patient outcome beyond either alteration alone. By systematically integrating survival associations with pairwise combinations of recurrent copy-number alterations and frequently mutated driver genes, we identified 57 PGIs. These PGIs refined prognostic stratification and were linked to distinct transcriptomic programs representing immune-response, epithelial-mesenchymal transition, and proliferation-related themes. Gene-level mapping highlighted dosage-sensitive candidate genes within recurrent copy-number regions, and gene essentiality profiles supported subsets of PGI-derived gene pairs. Two PGIs were validated in independent datasets. Together, these results establish a framework for identifying prognostic combinations of genomic alterations and connecting them to pathway programs, candidate genes, and functional dependencies in human tumors.

## Background

Cancer genomes harbor somatic mutations, copy-number alterations (CNAs), and broader chromosomal abnormalities that shape tumor behavior, therapeutic response, and clinical outcome. Genomic profiling is now widely used both to understand tumor biology and to improve prognostication and treatment selection for individual patients (van Putten *et al*, 2026; ICGC/TCGA Pan-Cancer Analysis of Whole Genomes Consortium, 2020; AACR Project GENIE Consortium, 2017).

However, clinically actionable cancer biomarkers are still largely defined by alterations in individual genes or by broad genomic signatures, rather than by combinations of genomic events (Suehnholz *et al*, 2024). This single-feature perspective may miss clinically meaningful subsets of tumors defined by combinations of alterations that together shape disease behavior, especially as genomic alterations are highly heterogeneous across patients and cancer types. Furthermore, genome-wide properties such as CNA burden and tumor mutational burden are also linked to tumor phenotypes, clinical outcome, and therapy response (Samstein *et al*, 2019; Hieronymus *et al*, 2018; Taylor *et al*, 2018). This further emphasizes the need for prognostic models that go beyond isolated genomic alterations to consider combinations of events within the broader genomic context of each tumor.

Combinations of genomic alterations may be especially informative because malignant phenotypes often emerge from cooperating events rather than single alterations acting alone. Consistent with this view, primary cancer genomes typically harbor around five driver mutations and frequently converge onto molecular pathways and interaction networks (ICGC/TCGA Pan-Cancer Analysis of Whole Genomes Consortium, 2020; Reyna *et al*, 2020), indicating that tumor behavior is shaped by combinations of genomic lesions rather than single events. In model organisms, systematic genetic-interaction mapping has revealed pathway structure, functional redundancy, and compensatory relationships that are not apparent from single-gene analyses (Costanzo *et al*, 2016; Lehner *et al*, 2006). In human cancer cell lines, complementary insights have come from large-scale perturbation screens and dependency maps that link genetic vulnerabilities to molecular context, including more recent combinatorial CRISPR screens that directly assay pairwise genetic interactions (Tsherniak *et al*, 2017; Hart *et al*, 2015; Shen *et al*, 2017; Gonatopoulos-Pournatzis *et al*, 2020). These experimental frameworks have been influential for discovering synthetic-lethal and other context-dependent genetic dependencies in cancer (O’Neil *et al*, 2017). Such approaches remain largely limited to experimental cancer systems, and linking combinatorial functional relationships to clinically annotated patient genomics cohorts remains a challenge. Addressing this gap could help reveal how combinations of genomic alterations shape tumor biology in patients, and identify prognostic biomarkers and candidate therapeutic vulnerabilities.

Despite growing recognition that tumor phenotypes are shaped by combinations of genomic events, systematic methods to identify clinically relevant alteration pairs directly from patient cohorts remain limited. Studies of co-occurring driver alterations have shown that specific combinations of genomic events can define biologically and clinically distinct tumor subgroups, particularly in oncogene-defined cancers, as exemplified by *KRAS*-mutant lung adenocarcinoma with *CDKN2A/B* inactivation (Skoulidis *et al*, 2015). Broadly, patterns of co-occurrence and mutual exclusivity among somatic alterations can reveal positive selection, pathway cooperation, redundancy, and subtype structure across cancer cohorts (El Tekle *et al*, 2021; Mina *et al*, 2017). Pairwise interaction frameworks have been used to relate joint molecular states to survival in cancer genomics cohorts, although mostly at the transcriptomic level (Magen *et al*, 2019). However, most cancer biomarker studies still emphasize single alterations or broad molecular classes, whereas most functional interaction studies are based on experimental systems such as cell lines and perturbational screens. Thus, approaches that systematically identify prognostic genomic alteration pairs directly from patient cohorts and connect them to matched transcriptomes and functional evidence remain limited.

To address this challenge, we analyzed genomic and clinical data from nearly 10,000 primary tumors across major cancer types to identify prognostic genomic interactions (PGIs). Building on our recently developed PACIFIC framework for survival-based interaction analysis (Bayati *et al*, 2025), we systematically evaluated pairwise combinations of recurrent CNAs and frequently mutated cancer driver genes for associations with patient outcome. We used matched transcriptomes and external functional resources to understand the biological context of PGIs and followed selected signals in independent clinical datasets. Together, this study provides a framework for discovering prognostic genomic alteration pairs and linking them to pathway programs, candidate genes, and functional evidence in human tumors.

## Results

### Systematic discovery of prognostic genomic interactions across cancer types

We defined a prognostic genomic interaction (PGI) as a pair of genomic features whose joint alteration status was associated with patient outcome beyond either feature alone, after adjusting for baseline covariates. We then modeled survival associations of genomic feature pairs using the PACIFIC framework (Bayati *et al*, 2025) separately within each of 24 cancer types spanning 8,359 tumors in total (**Table S1**). PACIFIC is an elastic net regularized survival analysis framework that repeatedly evaluates pairwise feature combinations across subsamples of the cohort and identifies robust interactions whose prognostic signals exceed the corresponding single features. Here, we evaluated all pairwise combinations of recurrent CNAs and frequently mutated cancer driver genes from TCGA studies (Bailey *et al*, 2018; Taylor *et al*, 2018). Because chromosomal instability and elevated CNA burden have been associated with more aggressive tumors and adverse clinical outcomes (Al-Rawi *et al*, 2024; Taylor *et al*, 2018), we included overall CNA burden as a covariate in all survival models, together with baseline clinical covariates of patient age, sex, tumor stage and grade as available.

PACIFIC identified 57 PGIs across 15 cancer types (**Figure 1A, Table S2**). Most PGIs were defined by pairs of CNAs (41/57, 72%), whereas the remainder involved small mutations in established driver genes, including mutation-CNA and mutation-mutation pairs (16/57, 28%). About half of all PGIs (29/57) were associated with worse prognosis. PGIs were distributed across cancer types, with most observed in lung adenocarcinoma (LUAD), colon adenocarcinoma (COAD), bladder urothelial carcinoma (BLCA), and IDH-wildtype glioblastoma (GBM), each contributing more than five PGIs to the final catalog.

**Figure 1.**
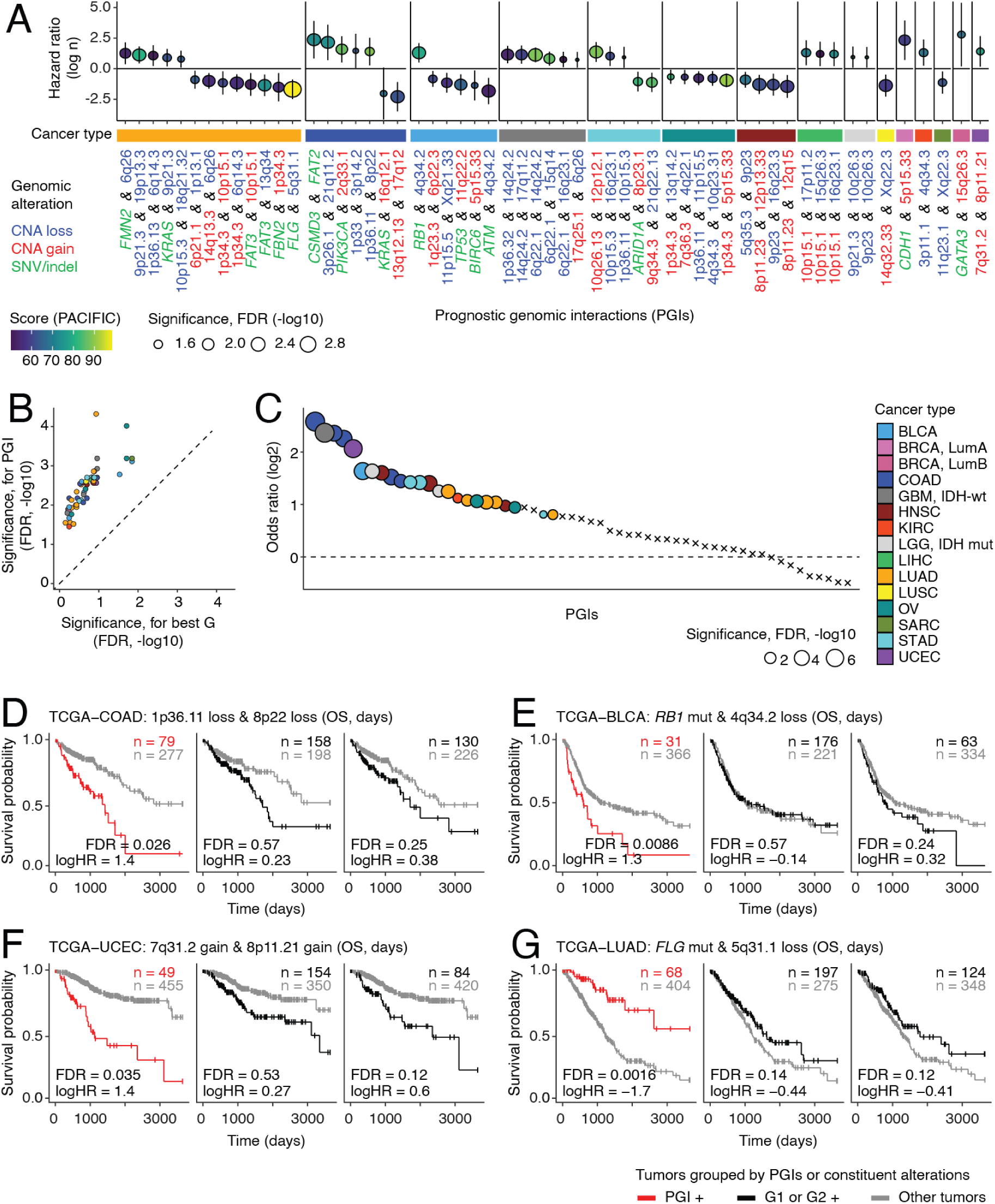
Prognostic genomic interactions (PGIs) discovered by survival analysis. **(A)** Catalog of 57 PGIs identified in survival analyses of individual cancer types, grouped by the cancer type in which each PGI was discovered. The x-axis lists PGIs and the y-axis shows the log hazard ratio (HR) from Cox proportional hazards (CoxPH) models. Models were adjusted for baseline clinical covariates and CNA burden, and the PGI-based models were also adjusted for the two constituent genomic features G1 and G2. Point size and color indicate the FDR-adjusted P-value and the importance score from PACIFIC, respectively. X-axis labels are colored by genomic fearture class: small mutation in driver gene (green), copy-number loss (blue), and copy-number gain (red). **(B)** PGIs showed stronger association with patient survival than either constituent genomic feature alone. Each point represents a PGI (colored by cancer type). The x-axis shows the stronger (smaller) FDR-adjusted P-value of either G1 or G2, derived from CoxPH models adjusted for baseline clinical covariates. The y-axis shows the FDR-adjusted P-value for the PGI term from CoxPH models adjusted for baseline clinical covariates and G1 and G2. **(C)** Co-occurrence of the two genomic features (G1, G2) comprising each PGI within the corresponding cohort, assessed by two-tailed Fisher’s exact tests. Points are colored by cancer type and sized by FDR-adjusted P-value. Crosses indicate non-significant associations (FDR > 0.05). The y-axis shows the log2 odds ratio. (**D-G)** Examples of PGIs and their associations with patient survival. Kaplan-Meier (KM) curves compare survival for PGI-positive versus PGI-negative patients (left of each panel). As controls, survival stratified by each constituent genomic feature alone (G1 or G2) is shown (middle and right of each panel). **(D)** Co-loss of 1p36.11 and 8p22 was associated with worse prognosis in colon adenocarcinoma (COAD). **(E)** *RB1* mutation with 4q34.2 loss was associated with worse prognosis in bladder urothelial carcinoma (BLCA). **(F)** Co-gain of 7q31.2 and 8p11.21 was associated with worse prognosis in uterine corpus endometrial carcinoma (UCEC). **(G)** *FLG* mutation with 5q31.1 loss was associated with better prognosis in lung adenocarcinoma (LUAD). Reported FDR-adjusted P-values and HRs were derived from CoxPH models adjusted for baseline clinical covariates, and for PGI models, G1 and G2 were additionally included as covariates.

To quantify the added prognostic value of paired genomic alterations, we compared each PGI with each of its two underlying genomic alterations. PGIs showed stronger survival associations than either alteration alone, with systematically more significant adjusted P-values than single-feature associations (**Figure 1B**). This pattern is consistent with the PGI modeling framework and confirms that combinations of alterations captured prognostic stratification beyond single genomic alterations and baseline covariates.

We asked whether the two genomic alterations comprising each PGI tended to co-occur within the corresponding tumor cohort more often than expected by chance. Overall, genomic alteration pairs in 24 of 57 PGIs showed significant co-occurrence (Fisher’s exact test; FDR < 0.05; **Figure 1C**). Such co-occurrence is consistent with non-random coupling of alterations within tumors, which may reflect joint positive selection, shared genomic context, or the characteristics of molecular cancer subtypes (El Tekle *et al*, 2021). Notably, most PGIs (33 of 57) did not show significant co-occurrence at the cohort level, indicating that PGI discovery was not restricted to highly recurrent co-alteration patterns but also captured less frequent co-alteration states associated with distinct outcomes.

We examined representative examples of PGIs (**Figure 1D-G**). Co-loss of the chromosomal regions 1p36.11 and 8p22 in colorectal cancer defined a subset of tumors with significantly worse prognosis (**Figure 1E**). At 1p36.11, *ARID1A* encodes a SWI/SNF chromatin-remodeling factor, and *ARID1A* loss has been shown to disrupt enhancer-mediated transcriptional control and promote colon tumorigenesis (Mathur *et al*, 2017). At 8p22, *DLC1* encodes a RhoGAP tumor suppressor, and restoration of *DLC1* has been shown to suppress cancer cell growth, invasion, and metastatic behavior, including in colorectal cancer models (Goodison *et al*, 2005; Wang *et al*, 2014). The combined loss of 1p36.11 and 8p22 is therefore compatible with joint suppression of chromatin-regulatory and invasion-suppressive programs, providing a plausible biological basis for the poor-prognosis subgroup identified in colon adenocarcinoma.

Additional PGIs further illustrated the diversity of genomic interactions across tumor types. In bladder urothelial carcinoma, *RB1* mutation together with 4q34.2 loss was associated with worse overall survival (**Figure 1D**), consistent with prior studies linking *RB1* disruption to aggressive muscle-invasive and neuroendocrine-like bladder cancer (Robertson *et al*, 2018). In uterine corpus endometrial carcinoma, co-gain of 7q31.2 and 8p11.21 also marked a poor-prognosis subgroup (**Figure 1F**). These loci harbor plausible oncogenic candidates, including the established oncogene *MET* at 7q31.2 (Comoglio *et al*, 2018) and the chromatin modifier and potential therapeutic target *KAT6A* at 8p11.21 (Mukohara *et al*, 2024; Turner-Ivey *et al*, 2014), suggesting that combined activation of growth-factor signaling and chromatin-regulatory programs may contribute to adverse outcome. In contrast, not all PGIs were associated with poor prognosis: in LUAD, *FLG* mutation together with 5q31.1 loss identified a subgroup with improved survival (**Figure 1G**), illustrating that PGIs captured both unfavorable and favorable co-alteration states across cancers.

Collectively, these results show that clinically relevant prognostic signals can be found from pairs of genomic alterations, even when the underlying events are not strongly co-occurring within a cohort. This prompted us to examine whether PGIs were linked to transcriptomic and pathway-level programs.

### PGIs cluster into immune, EMT, and proliferation pathway programs

To investigate whether PGIs were associated with distinct transcriptional consequences, we analyzed matched tumor transcriptomes for all 57 PGIs using differential expression models that tested the joint PGI effect while accounting for the individual effects of the two underlying genomic features. In several cases, PGIs were associated with extensive transcriptional remodeling, including three PGIs linked to differential expression of more than a thousand genes (**Figure 2A**). PGIs were often associated with expression changes in known cancer-relevant genes, indicating that co-alteration states were linked to biologically meaningful transcriptomic programs rather than only broad shifts in gene expression.

**Figure 2.**
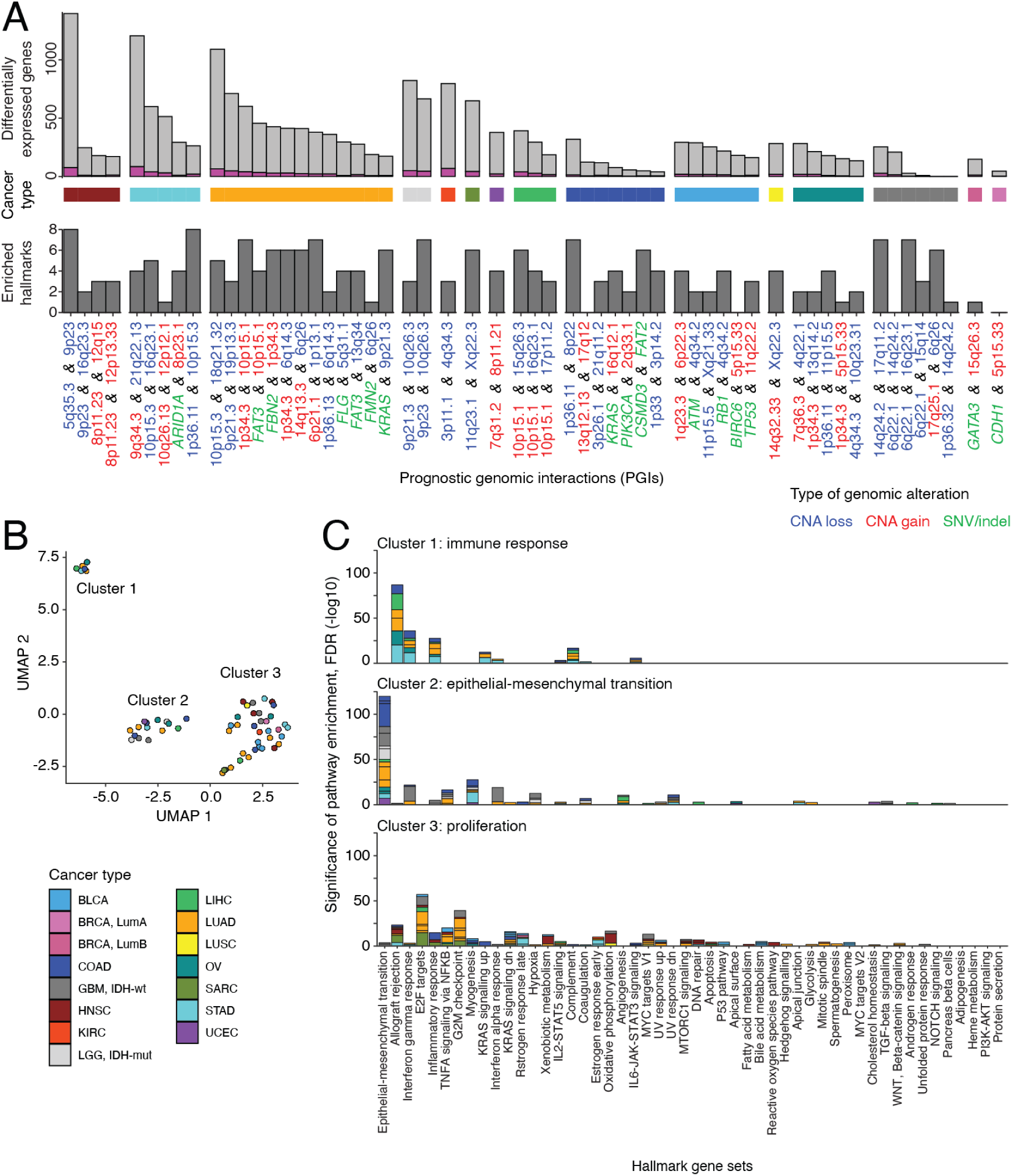
Transcriptome-derived pathway signatures of prognostic genomic interactions (PGIs). **(A)** Differential expression and pathway enrichment analyses for PGIs. For each PGI, we tested the PGI term for association with gene expression while accounting for the main effects of the two constituent genomic features (see Methods). The top panel shows the number of genes with significant PGI-associated differential expression (FDR < 0.05), for each PGI (x-axis). Among these, cancer-relevant genes are highlighted in purple. The bottom panel shows the number of significantly enriched Hallmark gene sets based on the 50 MSigDB database identified by ActivePathways using PGI-associated differential expression results (Holm family-wise error rate (FWER) < 0.05). The plot is faceted horizontally by the cancer type in which each PGI was identified. X-axis labels are colored by feature class: gene mutation (green), copy-number loss (blue), and copy-number gain (red). **(B)** Dimensionality reduction and clustering of PGIs based on enrichment profiles of Hallmark gene sets from MSigDB. The UMAP algorithm was used for visualization and the k-means algorithm for clustering. **(C)** Hallmark gene sets enriched across PGIs within each cluster of three clusters (immune response, epithelial-mesenchymal transition, and proliferation). Bars summarize Hallmark gene sets significantly enriched across PGIs in each of the three clusters (FWER < 0.05) in which each bar segment represents a given PGI. Colors in panels B and C indicate the cancer type associated with each PGI.

We next asked whether these PGI-associated transcriptional changes converged onto shared biological pathways. Most PGIs (54/57) were associated with Hallmark signatures from the MSigDB database (Liberzon *et al*, 2015) based on pathway enrichment analysis (family-wise error rate (FWER) < 0.05 from ActivePathways (Paczkowska *et al*, 2020)) (**Figure 2A, Table S3**). Several PGIs showed especially strong pathway-level signals. For example, the PGI involving co-loss of 10p15.3 and 18q21.32 was associated with worse prognosis in LUAD and differential expression of 1090 genes. The deleted 18q21 region contains *SMAD4*, a central mediator of canonical TGF-β signaling, and this PGI was accompanied by a strong enrichment of EMT-related processes in our pathway analysis (FWER = 3.3 x 10^-15^), consistent with invasive and metastasis-associated programs in LUAD (Lee *et al*, 2024; Derynck *et al*, 2021). Thus, some PGIs corresponded to coherent pathway-level programs rather than isolated gene-expression changes.

To compare pathway profiles across PGIs, we represented each PGI by its Hallmark enrichment profile and performed dimensionality reduction and clustering. This analysis separated PGIs into three broad groups defined by dominant pathway themes: an immune-response cluster, an EMT cluster, and a proliferation cluster (**Figure 2B,C**). The immune-response cluster was characterized by inflammatory and interferon-related signatures, the EMT cluster by mesenchymal transition, extracellular matrix, adhesion, and related stromal-remodeling programs, and the proliferation cluster by cell-cycle, mitotic, and growth-associated pathways. These clusters were not confined to a single cancer type, indicating that distinct PGIs arising in different tumor contexts can converge on similar programs defined by transcriptional and pathway-level signatures.

Together, these analyses showed that PGIs were not only prognostic but also biologically interpretable. By linking co-alteration states to recurrent pathway programs, transcriptomic analysis provided a functional framework for organizing PGIs beyond survival associations alone and highlighted immune, EMT, and proliferative states as major axes of PGI-associated tumor biology.

### Candidate dosage-sensitive target genes for CNA features in PGIs

Because the CNA features in PGIs spanned recurrent chromosomal regions that could affect multiple genes, we next mapped these to candidate dosage-sensitive target genes by identifying significant, CNA-concordant expression changes between PGI-positive and PGI-negative tumors (DESeq2, FDR < 0.05). We then enumerated the resulting gene-gene pairs at two levels of stringency. Restricting candidate dosage-sensitive genes to known cancer-relevant genes yielded 385 gene pairs across 20 PGIs (**Figure 3A,B**), whereas the full analysis identified 38,189 gene pairs across 39 PGIs overall (**Table S4**). Thus, restricting the analysis to known cancer-relevant genes substantially reduced the candidate space while preserving a focused set of interpretable gene-level hypotheses. Among the 385 pairs of cancer-relevant genes, we identified ten gene pairs across seven PGIs comprising pairs of CNAs and recurrently mutated genes, including major driver genes *RB1*, *TP53*, and *KRAS* (**Figure 3B**). These results suggest that at least a subset of PGIs may reflect coordinated dosage effects on biologically important genes affected by recurrent CNAs.

**Figure 3.**
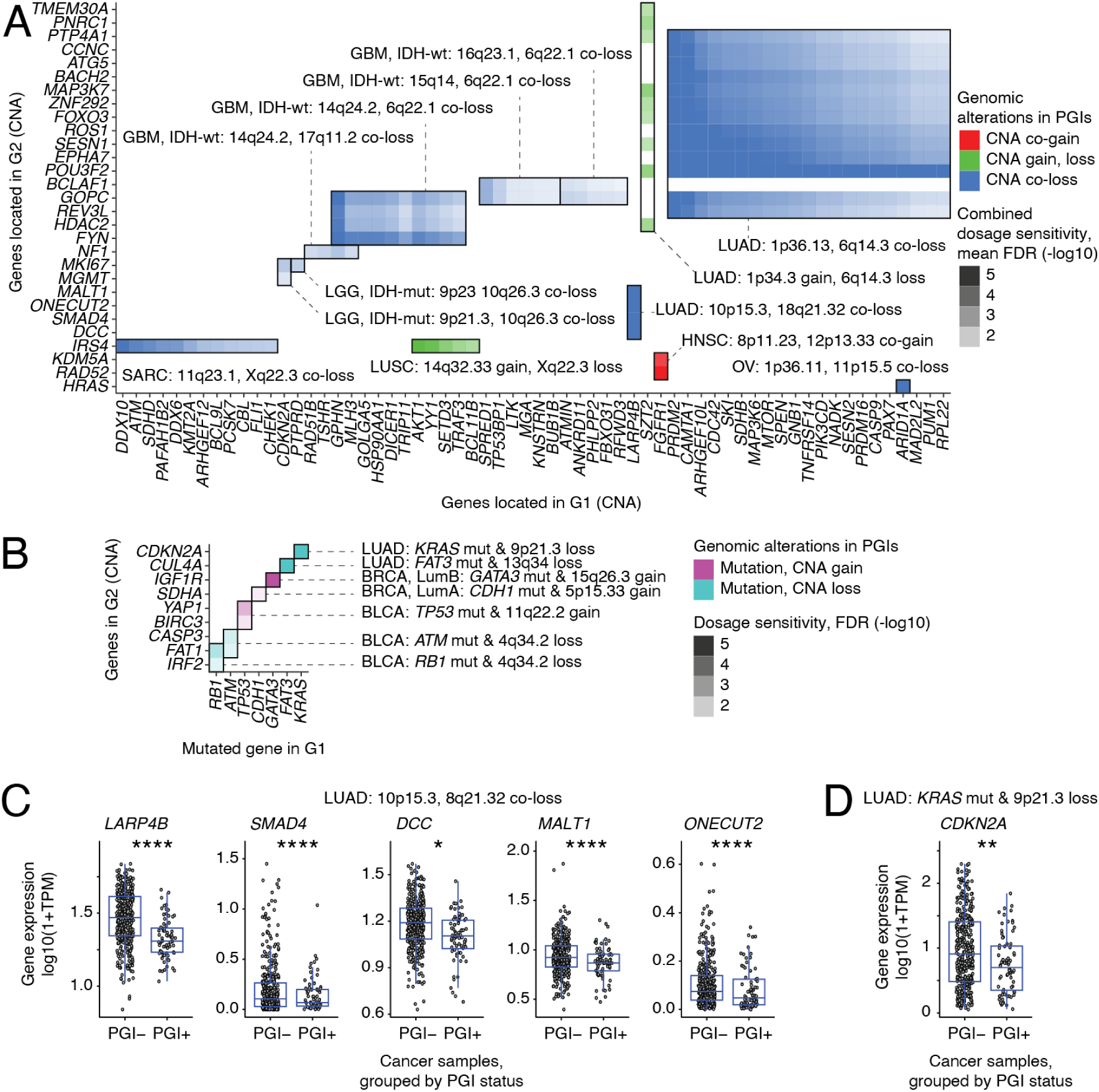
Candidate dosage-sensitive target genes for PGIs involving recurrent copy number alterations (CNAs). **(A)** Candidate gene-level targets for PGIs involving pairs of CNAs (denoted as G1, G2), with co-located genes shown on the X and Y axes, respectively. Each rectangle indicated by black borders corresponds to one PGI and represents a pair of candidate target genes mapped from the two constituent CNAs. Candidate genes were restricted to known cancer genes within each CNA interval that showed differential expression concordant with CNAs and reflecting dosage sensitivity (DESeq2, FDR < 0.05). Tile colors indicate the direction of differential expression for both genes. Color gradient indicates the strength of differential expression, quantified as the mean significance for the two genes, where negative log10-transformed FDR values were derived from differential gene expression analyses. **(B)** Candidate gene-level targets for mutation-CNA PGIs. Each rectangle indicated by black borders corresponds to a PGI involving a pair of a CNA (Y axis) and mutated cancer gene (X axis). Candidate genes in CNAs are derived similarly to panel A. **(C)** Dosage sensitivity of cancer-relevant genes for the PGI involving co-loss of 10p15.3 and 18q21.32 in LUAD. **(D)** Dosage sensitivity of *CDKN2A* for the PGI involving KRAS mutation and 9p21.3 loss in LUAD. Boxplots expression of candidate genes in PGI-positive versus PGI-negative tumors in corresponding cancer types. Expression is quantified as log10-transformed TPM values. Asterisks indicate significance of differential expression (* P < 0.05; ** P < 0.01; *** P < 0.001; **** < 10^-4^).

An illustrative CNA-CNA example was the co-loss of 10p15.3 and 18q21.32 in LUAD, which was also associated with a strong transcriptomic and pathway signals described above. PGI-positive tumors showed reduced expression of the TGF-β regulator *SMAD4*, together with *MALT1*, *ONECUT2*, and *DCC* at 18q21.32 and *LARP4B* at 10p15.3, consistent with dosage-sensitive attenuation of multiple genes across both deleted loci (**Figure 3C**). This pattern suggests coordinated dosage loss across multiple candidate genes.

For a mutation-CNA example, we highlighted the PGI defined by *KRAS* mutation and 9p21.3 loss in LUAD. This PGI mapped to the gene pair *KRAS*-*CDKN2A*, and *CDKN2A* expression was reduced in PGI-positive tumors relative to PGI-negative tumors, consistent with deletion of the 9p21.3 locus (**Figure 3D**). This result supports the biological plausibility of the mapped gene pair and illustrates how mutation-CNA PGIs can connect recurrent driver mutations to dosage-sensitive genes in altered chromosomal regions.

Overall, the predicted gene-level resolution varied across PGIs. In some cases, CNA features converged on a small number of interpretable candidate genes, whereas in others they pointed to broader multi-gene consequences of recurrent genomic loss or gain. Accordingly, dosage-sensitivity mapping provided a principled framework for prioritizing candidate genes and translating region-level PGIs into testable gene- and locus-level hypotheses arising from prognostic combinations of genomic alterations.

### Co-essential networks and synthetic-lethal annotations of PGI-derived gene pairs

We next asked whether dosage-sensitive candidate gene pairs from PGIs were also supported by functional evidence for genetic interactions in cancer model systems, focusing on gene co-essentiality and curated synthetic-lethal relationships. Using CRISPR-Cas9 dependency scores across 1186 cell lines profiled in the DepMap project (Tsherniak *et al*, 2017), we tested whether essential phenotypes for the two genes in each PGI-derived pair co-occurred more often than expected by chance.

This analysis identified 103 significant co-essential gene pairs involving 77 genes and six PGIs (FDR < 0.05; **Figure 4A**). Examining the co-essentiality network, we found that gene pairs clustered into a few major connected components representing core cellular processes, including centrosome and microtubule organization (*TUBE1*), mitochondrial translation and tRNA biology (*RARS2*, *TRIT1*), DNA replication (*ORC3*), and transcriptional regulation (*MED23*). Well-known cancer genes were also present, including *SMAD4*, *ARID1A*, *RUNX3*, *DICER1*, *HDAC2*, and *SDHB*, indicating that PGI-derived gene pairs were not limited to generic cellular fitness genes but also encompassed genes with established relevance to tumor biology.

**Figure 4.**
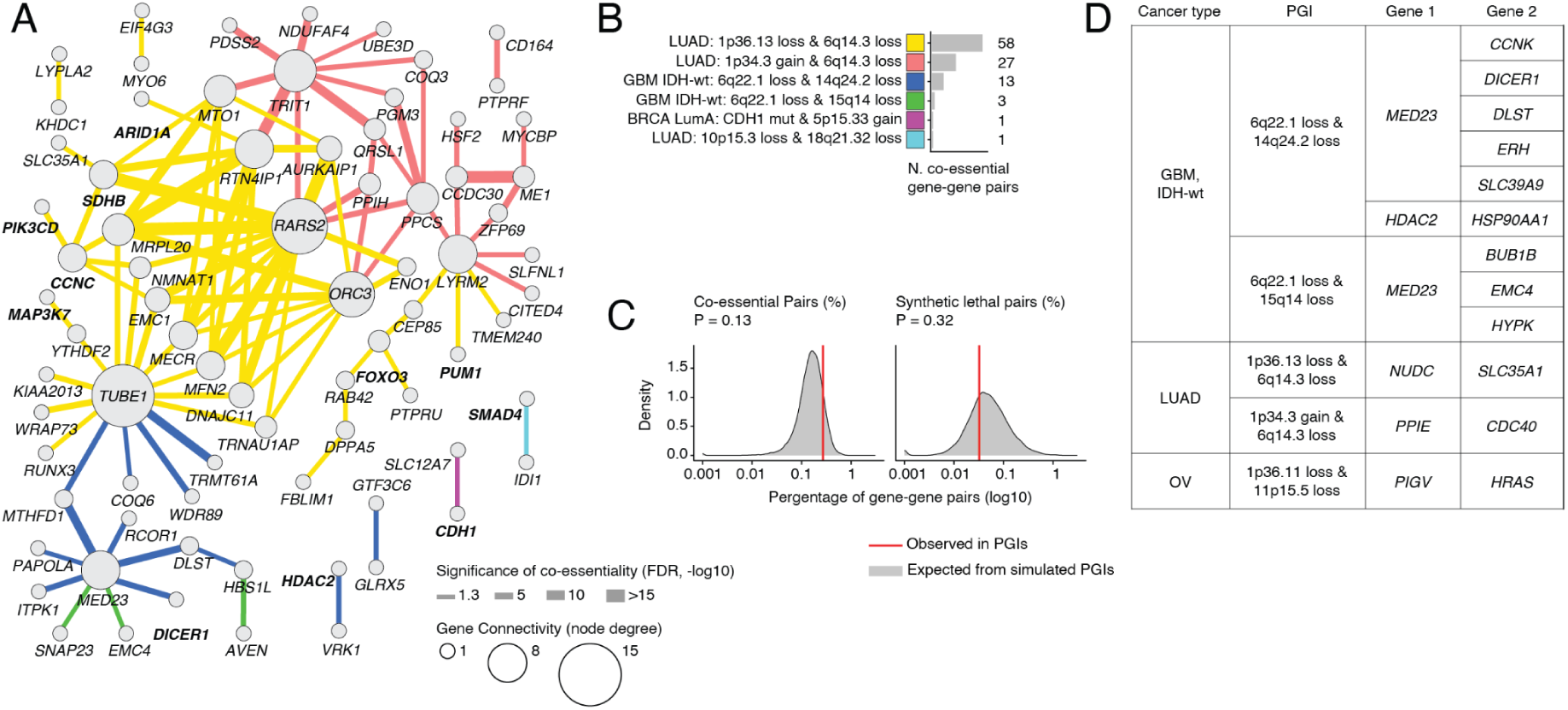
Overlap of PGI-derived gene-gene pairs with co-essentiality and curated synthetic-lethal interactions. **(A)** Network of co-essential relationships among genes implicated by PGIs, based on DepMap CRISPR-Cas9 gene-dependency data across 1186 cell lines. Nodes represent genes and node size indicates degree (number of connections). Edges indicate significant co-essential PGI-derived gene pairs identified from Chronos dependency scores from PGI-derived gene-gene pairs (one-tailed Fisher’s exact test; FDR < 0.05). Known cancer-related genes are highlighted in boldface. **(B)** Number of significant co-essential gene pairs in panel A attributable to each PGI. **(C)** Permutation analysis of co-essential and synthetic-lethal gene pairs derived from simulated PGIs with matched dosage-sensitivity analysis. Density plots show the expected fraction of co-essential or synthetic-lethal gene pairs across 10,000 simulated PGI sets matched by genomic alteration class and cancer type. Red vertical lines indicate the observed fractions from the true PGIs. Empirical P-values are shown, and error bars indicate 95% intervals for the expected fractions. **(D)** PGI-derived gene pairs annotated as synthetic-lethal (SL) in SynLethDB.

Although co-essential pairs were detected across multiple PGIs, most (85/103) were derived from two PGIs identified in LUAD (**Figure 4B**). To test whether this level of co-essentiality exceeded expectation by chance, we performed a stringent custom permutation analysis by constructing simulated PGIs from randomly sampled genomic feature pairs matched by cancer type and genomic event class (CNA-CNA or mutation-CNA), followed by equivalent testing of dosage sensitivity. PGI-derived gene pairs showed a higher proportion of co-essential relationships than matched random pairs, but this difference was not statistically significant (0.27% observed vs. 0.17% expected; one-tailed P = 0.13; **Figure 4C**).

We next compared PGI-derived gene pairs with curated synthetic-lethal relationships from the SynLethDB database (Guo *et al*, 2016). This analysis identified 12 synthetic-lethal gene pairs derived from five PGIs (**Figure 4D**). Nine gene pairs arose from two PGIs identified in IDH-wildtype GBM, defined by co-loss of 6q22.1 and 14q24.2 or co-loss of 6q22.1 and 15q14, respectively, and involved the Mediator complex subunit *MED23*, a core transcriptional regulator with established cancer relevance (Yang *et al*, 2012), indicating recurrent synthetic-lethal roles for this gene in our analysis. Additional annotated interactions included *HDAC2-HSP90AA1*, a synthetic-lethal pair from SynthLethDB that has potential therapeutic relevance given prior evidence for synergistic anticancer effects of combined HDAC and HSP90 inhibition and the development of dual inhibitors (George *et al*, 2005; Wu *et al*, 2021). A few established cancer-relevant genes such as *DICER1*, *BUB1B*, and *HRAS* were also identified. We adapted the permutation framework to test whether the overlap between PGI-derived gene pairs and curated synthetic-lethal interactions differed from the expected distribution. PGI-derived gene pairs did not show significant enrichment or depletion relative to chance (P = 0.32). Together, co-essentiality and curated synthetic-lethal annotations provided functional support for a subset of PGI-derived gene pairs, indicating that some PGIs can be linked to plausible biological mechanisms beyond their association with patient outcome.

### Clinical validation and multi-omics interpretation of two PGIs in lung adenocarcinoma and glioblastoma

We attempted external clinical validation of PGIs using independent cancer genomics datasets available through the cBioPortal database (Cerami *et al*, 2012) and obtained support for two PGIs: *KRAS* mutation with 9p21.3 loss in LUAD and co-loss of 6q22.1 and 15q14 in IDH-wildtype GBM (**Figure 5A-D**). In both cases, the joint alteration status identified patient subsets with worse overall survival in the TCGA discovery cohorts and retained prognostic significance in independent genomics datasets from the GENIE project (AACR Project GENIE Consortium, 2017). Because the discovery of PGIs involved genome-wide CNA features whereas the validation data were generated by targeted panel sequencing, validation of CNA-based features used selected genes within the corresponding regions. To interpret the two validated PGIs functionally, we examined their transcriptomic signatures in TCGA and mapped the corresponding CNA features to candidate genes within the affected loci. In LUAD, the PGI combining *KRAS* mutation with 9p21.3 loss was accompanied by a broad differential expression signature of 174 differentially expressed genes (FDR < 0.05, **Figure 5E, Table S5**), consistent with the distinct biological subgroup of *KRAS*-mutant LUAD with *CDKN2A/B* inactivation (Skoulidis *et al*, 2015). In the validation dataset, 9p21.3 loss was represented by *CDKN2A*, providing a direct bridge between the discovery signal from genomic regions and the gene-level signals from targeted panel data.

**Figure 5.**
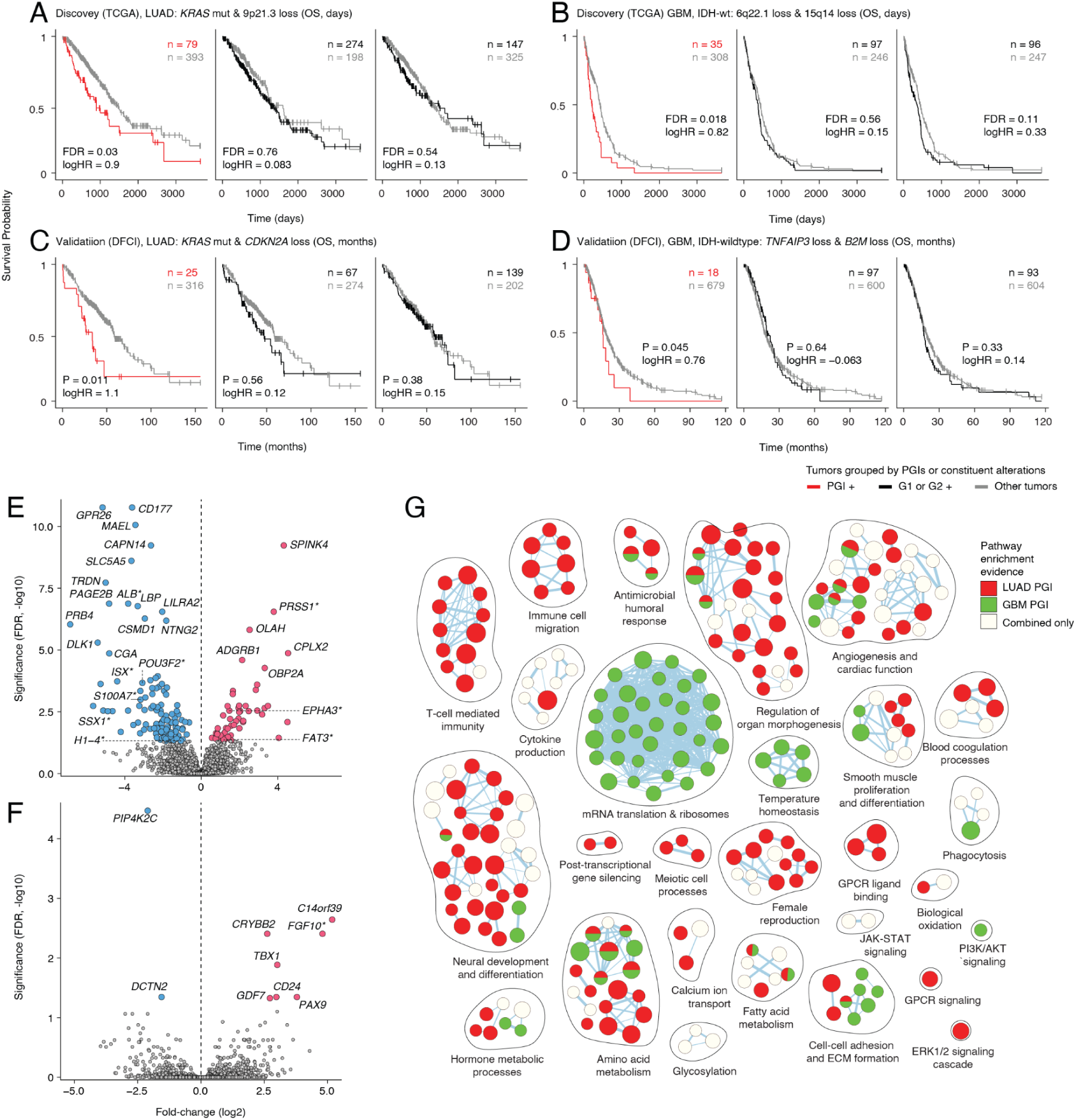
Two PGIs in lung adenocarcinoma (LUAD) and IDH-wild-type glioblastoma (GBM) associate with worse survival and distinct transcriptomic programs. (A-D) Association of two PGIs with overall survival (OS) in independent cohorts. **(A,B)** Discovery in TCGA: (A) *KRAS* mutation with 9p21.3 loss (*CDKN2A* locus) in LUAD, and **(B)** co-loss of 6p22.1 and 15q14 in IDH-wildtype GBM. **(C,D)** External validation in the Dana-Farber Cancer Institute (DFCI) subset of Project GENIE for LUAD **(C)** and IDH-wildtype GBM **(D)**. In each panel, Kaplan-Meier curves (left) compare PGI-positive versus PGI-negative samples. Control analyses stratified by either constituent genomic feature alone are shown (G1, middle; G2 right). Sample sizes (n) are indicated. P-values and hazard ratios (HRs) were obtained from Cox proportional hazards (CoxPH) models adjusted for baseline covariates (age and sex in all cohorts; tumor stage added for LUAD; CNA burden added for TCGA). PGI models additionally included for the constituent genomic features G1 and G2. **(E,F)** Differential expression analysis of the two PGIs in LUAD **(E)** and IDH-wildtype GBM **(F)**. In the volcano plots, each point represents a gene and points are colored by significant up- or downregulation (DESeq2, FDR < 0.05). Log2 fold change and FDR-adjusted P-value are shown on the x- and y-axes, respectively. Significant cancer-relevant genes are highlighted with asterisks in both panels; additionally, other genes meeting the more stringent thresholds are labelled (FDR < 0.001 in panel E; FDR < 0.05 in panel F). **(G)** Integrative pathway enrichment analysis of the gene signatures of the two PGIs using Reactome pathways and Gene Ontology biological processes. Circles represent significantly enriched pathways or processes (ActivePathways, FDR < 0.05). Pathways with substantial gene overlap are grouped into subnetworks and annotated with functional themes. Node size is proportional to pathway gene set size, and node color indicates enrichment supported by either of the two PGIs or both.

In the PGI found in IDH-wildtype GBM, the deleted regions 6q22.1 and 15q14 were represented by the co-located candidate genes *TNFAIP3* and *B2M*. *TNFAIP3* encodes A20, a negative regulator of NF-κB and inflammatory signaling, and recurrent inactivation of *TNFAIP3* has implicated it as a tumor suppressor in several lymphoid malignancies (Hertens & van Loo, 2024; Honma *et al*, 2009). *B2M* encodes β2-microglobulin, an essential component of MHC class I antigen presentation, whose alterations have been associated with impaired response to cancer immunotherapy (Sade-Feldman *et al*, 2017; Yang *et al*, 2023), consistent with immune evasion through defective antigen presentation. Transcriptomic analysis of the 6q22.1/15q14 co-deletion PGI revealed a more modest signature, with nine differentially expressed genes (FDR < 0.05, **Figure 5F, Table S5**).

Although the two validated PGIs differed in the magnitude of their gene-level expression signatures, both could be placed in a common pathway-level context. Integrative pathway enrichment analysis across the two PGIs identified 293 significantly enriched pathways and processes (ActivePathways; FDR < 0.05), which clustered into coherent functional themes and collectively captured signals from both PGIs (**Figure 5G, Table S6**). The major functional themes involved immune processes, translational and ribosome-related programs, cell adhesion and extracellular matrix organization, angiogenesis, and developmental functions, including programs related to neural differentiation. Together, these patterns indicate that the validated PGIs map onto coordinated tumor states involving microenvironmental interaction, biosynthetic activity, and tissue remodeling. Thus, pathway-level interpretation connected the two validated PGIs to broader tumor biology beyond individual differentially expressed genes.

## Discussion

Our analysis shows that prognostic information in cancer genomes can emerge from combinations of genomic alterations rather than from single events considered in isolation. We identified PGIs in multiple cancer types whose joint alteration status refined patient stratification beyond either underlying feature alone, even after accounting for baseline covariates and overall CNA burden. These results suggest that clinically meaningful tumor states are often defined by co-alteration patterns embedded within a broader genomic context, extending current biomarker frameworks while remaining interpretable in relation to patient outcome.

Many PGIs could also be placed in a broader biological context through matched transcriptomes, gene-level mapping, and external functional resources. At the pathway level, PGIs converged on a limited number of recurrent programs, notably immune and microenvironmental signaling, EMT and tissue remodeling, and proliferative states, suggesting that diverse co-alteration patterns can map onto common axes of tumor biology. At a finer level, dosage-sensitivity mapping helped translate recurrent CNA features into candidate genes and testable locus-level hypotheses, including many established cancer-related genes whose expression changed concordantly with the underlying CNA events. Co-essentiality and curated synthetic-lethal annotations provided complementary functional support for a subset of PGI-derived gene pairs, indicating that some PGIs may reflect biologically coupled alterations with shared dependencies or candidate vulnerabilities. Taken together, these analyses provided a practical framework for linking PGIs to candidate genes, pathways, and systems-level hypotheses about tumor biology.

The two validated PGIs illustrate both the promise and the interpretive value of this framework. In LUAD, the prognostic interaction of *KRAS* and 9p21.3/*CDKN2A* recapitulates a biologically meaningful co-alteration pattern that has been linked to a distinct *KRAS*-mutant subgroup of LUAD (Skoulidis *et al*, 2015; Schuster *et al*, 2014). *CDKN2A* co-alterations are associated with worse outcomes in patients with *KRAS* G12C-mutant NSCLC treated with KRAS inhibitors (Negrao *et al*, 2023), highlighting the potential translational relevance of this PGI in the treatment setting. By contrast, the validated PGI in IDH-wildtype GBM, defined by co-loss of 6q22.1 and 15q14, represents a less-characterized prognostic interaction; however, the involvement of *TNFAIP3* and *B2M* points to a plausible link with inflammatory signaling and immune-related tumor biology. Together, these case studies show how PGI discovery in patient cohorts can recover both established and less-characterized co-alteration patterns with clinical and biological relevance.

Our findings position PGIs at the intersection of biomarker discovery and functional cancer genomics. Prior studies have shown that co-occurring driver alterations can define clinically distinct tumor subgroups, whereas large-scale perturbation screens have uncovered context-specific functional relationships in model systems. PGIs complement these approaches by identifying prognostic alteration pairs directly from clinically annotated patient cohorts and connecting them to transcriptomic programs, candidate genes, and complementary functional evidence. Thus, PGIs can highlight candidate pairs of genomic alterations and generate hypotheses about shared dependencies, pathway convergence, and vulnerabilities for downstream study.

Several limitations should be considered when interpreting these findings. First, PGIs are defined through associations with patient outcome and therefore do not by themselves establish direct genetic interaction, causal effects, or functional cooperation between the underlying alterations. Second, external validation was feasible for only a small subset of PGIs because of the limited availability of independent datasets, particularly where panel-based sequencing data required gene-level estimates of broad CNA regions. Third, mapping recurrent CNA features to candidate genes remains context-dependent and sometimes ambiguous, especially for larger regions in which multiple dosage-sensitive genes are located.

Fourth, transcriptomic and pathway analyses were performed at the cohort level and provide functional context rather than direct mechanistic proof. Finally, co-essentiality and curated synthetic-lethal resources remain incomplete and biased toward currently studied genes, model systems, and interaction types, so the absence of support in these resources should not be interpreted as evidence against biological relevance.

Taken together, our results suggest that survival-based analysis of genomic alteration pairs can complement existing approaches to cancer genomics by identifying clinically relevant combinations of genomic alterations with interpretable biological context. Conceptually, it also offers a way to study how networks of genes shape tumor phenotypes in human cohorts. Although the present study focused on overall survival, the same framework could be extended to other clinically relevant endpoints, including therapy response, disease progression, and treatment resistance. This may be especially informative in settings where combinations of alterations influence therapeutic sensitivity, as exemplified by the clinically established relationship between *BRCA1/2* deficiency and PARP inhibitor response (Bryant *et al*, 2005). Extending PGI analysis in this direction may help connect prognostic co-alteration patterns to predictive biomarkers and candidate therapeutic vulnerabilities.

## Methods

### Cancer samples

We obtained data for 10,960 primary tumors spanning 32 solid cancer types from the TCGA PanCancer Atlas (PanCanAtlas) project (Hoadley *et al*, 2018). Each case included matched somatic genomic data, bulk RNA-sequencing profiles, and clinical annotations, and each cancer type was analyzed independently. Genomic profiling in the TCGA project was performed in prior studies conducted under the U.S. Common Rule and local IRB requirements with written informed consent from all participants. All analyses in this study used publicly accessible datasets. For clinical outcome analyses, we used overall survival (OS) or progression-free survival (PFS) as defined in published TCGA endpoint guidelines (Liu *et al*, 2018). Samples lacking survival information were excluded. Survival time was truncated at 10 years and right-censored. Baseline covariates in survival models included age, sex, and overall CNA burden, and we also included tumor stage and grade when available for most samples within a cancer type. Samples missing the required covariates were excluded for those analyses.

### Genomics and transcriptomics data

Genomic alteration data comprised somatic single-nucleotide variants (SNVs) and small insertions/deletions (indels) from whole-exome sequencing, copy-number alterations (CNAs) measured using Affymetrix SNP 6.0 arrays, and transcriptomic profiles from bulk RNA-seq. RNA-seq data was downloaded from the Genomic Data Commons (https://portal.gdc.cancer.gov, 2023-01-25) and was represented as TPM values. We additionally incorporated a previously published per-sample CNA burden metric defined as the proportion of genomic bases deviating from baseline ploidy (Thorsson *et al*, 2018); samples lacking this metric were excluded. Breast tumors were stratified into five PAM50 molecular subtypes, and lower-grade glioma and GBM were stratified by mutation status of IDH genes (IDH-mutant vs IDH-wildtype). Cancer types with fewer than 100 samples or with infrequent survival events (<5%) were excluded. After applying these criteria, the final dataset consisted of 8,359 tumors across 24 cancer types: bladder urothelial carcinoma (BLCA); breast invasive carcinoma, PAM50 basal-like subtype (BRCA-Basal); breast invasive carcinoma, PAM50 luminal A subtype (BRCA-LumA); breast invasive carcinoma, PAM50 luminal B subtype (BRCA-LumB); cervical squamous cell carcinoma and endocervical adenocarcinoma (CESC); colon adenocarcinoma (COAD); esophageal carcinoma (ESCA); glioblastoma multiforme, IDH-wild-type (GBM-IDHWT); head and neck squamous cell carcinoma (HNSC); kidney renal clear cell carcinoma (KIRC); kidney renal papillary cell carcinoma (KIRP); brain lower grade glioma, IDH-mutant (LGG-IDHMut); liver hepatocellular carcinoma (LIHC); lung adenocarcinoma (LUAD); lung squamous cell carcinoma (LUSC); ovarian serous cystadenocarcinoma (OV); pancreatic adenocarcinoma (PAAD); pheochromocytoma and paraganglioma (PCPG); prostate adenocarcinoma (PRAD); rectum adenocarcinoma (READ); sarcoma (SARC); stomach adenocarcinoma (STAD); thyroid carcinoma (THCA); and uterine corpus endometrial carcinoma (UCEC) (**Table S1**).

### Driver mutations and recurrent CNAs

We considered two categories of genomic alterations. First, we analyzed somatic small mutations (SNVs and indels) in 633 known cancer driver genes in the intOGen database (Martínez-Jiménez *et al*, 2020) (release 2024-09-20). Only protein-coding mutations were included, and synonymous mutations were excluded. Second, we analyzed recurrent CNAs defined from TCGA PanCanAtlas copy-number profiles using GISTIC2 with standard parameters (false discovery rate < 0.25; confidence level 0.99) (Mermel *et al*, 2011). CNA peak calls defined in the GISTIC2 file all_lesions were obtained from the Broad GDAC Firehose repository (downloaded 2020-05-04). The number of recurrent CNAs varied across cancer types, ranging from two in THCA to 93 in UCEC (**Table S1**). Throughout this study, recurrent CNAs and small mutations in driver genes are collectively referred to as genomic features.

### Discovery of prognostic genomic interactions (PGIs)

We defined a prognostic genomic interaction (PGI) as a pair of genomic features whose joint alteration status was associated with patient outcome beyond the association of either feature alone, after adjusting for baseline covariates (patient age, sex, tumor stage and grade when available, and overall CNA burden). We used the PACIFIC pipeline (Bayati *et al*, 2025) to discover survival-associated PGIs within each cancer type. Candidate predictors included all genomic features (main effects) and all pairwise co-alteration indicators between genomic features. The number of PACIFIC iterations was set to 100,000, and default values for other parameters were used. To further compare each PGI with its two constituent genomic features, we performed Cox proportional hazards (CoxPH) survival analyses within each cancer type using the same baseline covariates. We assessed survival differences by PGI status and, as controls, by each genomic feature alone. Statistical significance of each survival association was assessed using a likelihood-ratio test (i.e., ANOVA on nested CoxPH models), in which the null model included baseline covariates only, and the alternative model included baseline covariates plus either the PGI term or a single genomic feature term, followed by multiple testing correction using FDR as defined in the PACIFIC method. The PGI model further included both constituent genomic features as covariates.

### Mapping CNAs to potential target genes and dosage sensitivity

Each CNA feature in our analysis corresponds to a recurrent genomic interval that is frequently gained or lost in the corresponding TCGA cohort. We mapped each CNA feature to a set of candidate protein-coding target genes using three criteria: (1) location of the gene within the CNA interval, (2) relevance to cancer based on curated databases, and (3) evidence of dosage sensitivity, with concordant differential gene expression with the CNA event in the corresponding cohort (upregulated with CNA gains; downregulated with CNA losses). Cancer-relevant genes were defined as the union of two established databases of cancer driver genes: the Cancer Gene Census list (CGC, v102; downloaded 2025-07-01) from the COSMIC database (Sondka *et al*, 2018) and the Cancer Gene List from the OncoKB database (Chakravarty *et al*, 2017) (downloaded 2025-07-21). To assess concordant expression changes, we performed differential expression analyses using the DESeq2 method (Love *et al*, 2014) for each PGI, with TCGA gene-level RNA-seq counts as input and and PGI status as the binary variable (i.e., PGI-positive vs all other tumors in the cohort). For each analysis, P-values were adjusted for multiple testing using the Benjamini-Hochberg FDR method (Benjamini & Hochberg, 1995).

### Gene co-essentiality analysis

We analysed gene co-essentiality using DepMap CRISPR-Cas9 gene dependency scores from the Chronos method (Tsherniak *et al*, 2017; Dempster *et al*, 2021) (public release 25Q3). For each gene pair, DepMap dependency scores were extracted across all available cell lines. Cell lines with missing values for either gene were excluded. Gene essentiality was defined using a fixed threshold of Chronos score < −0.5, classifying each gene in each cell line as essential or non-essential. Genes classified as commonly essential in the DepMap Gene Dependency Profile Summary (Chronos_Combined) were excluded from further analysis. Gene pairs in which either gene was essential in fewer than five cell lines were also excluded. Enrichments were assessed using a one-tailed Fisher’s exact test, followed by FDR correction for multiple testing.

### Permutation analysis of co-essential gene pairs

We performed a permutation analysis to test whether PGI-derived gene-gene pairs showed more co-essentiality than expected by chance. Starting from the full set of genomic feature pairs considered by PACIFIC, we applied the same sparsity filter used in the original analysis and then generated 10,000 random catalogs of feature pairs while preserving the number of detected true PGIs, the observed distribution across cancer types, and the distribution of pair classes (CNA-CNA, mutation-CNA), further excluding any pairs occurring on the same chromosome. For each simulated catalog, gene-gene pairs were derived using the same procedure as for the observed PGIs, including differential expression-based mapping of dosage-sensitive genes. For each run we then quantified the total number of derived pairs, the number of testable pairs in DepMap, the number of significant co-essential pairs (FDR < 0.05), and the proportion of significant co-essential pairs among tested pairs. The observed values were compared against the resulting null distributions to derive empirical P-values.

### Mapping PGIs to synthetic-lethal gene pairs

To determine whether PGIs correspond to known genetic interactions, we retrieved gene-gene interaction annotations from the SynLethDB database (Guo *et al*, 2016) (downloaded 2024-12-12), which catalogs five annotation types: synthetic-lethal (SL), non-synthetic lethal, synthetic dosage lethal, synthetic rescue, and synthetic dosage rescue. Because a gene pair can have multiple annotations due to evidence from multiple sources, we assigned each gene pair a single annotation using a weighted score based on annotation sources as described earlier (Wang *et al*, 2022; Guo *et al*, 2016), while excluding computationally predicted interactions. Next, we mapped each PGI to a set of candidate gene-gene interactions. Genomic features comprising small mutations (SNVs, indels) were mapped directly to their corresponding genes, whereas CNA features were mapped to candidate target genes within the CNA interval that showed concordant differential expression, as described above. We then assessed overlap between PGI-derived gene-gene interactions and SL pairs from SynLethDB. Statistical significance of SL gene pairs within the PGIs was also evaluated using a permutation test using the stringent process described above.

### External clinical validation of two PGIs using Project GENIE data

We reviewed cancer genomics datasets available in the cBioPortal resource (Cerami *et al*, 2012) to identify studies with sufficient sample size and the required genomic and clinical annotations to test PGIs for association with patient survival. We identified validation signals in the Dana-Farber Cancer Institute (DFCI) subset of Project GENIE (AACR Project GENIE Consortium, 2017), a targeted panel sequencing dataset focusing on known cancer genes. We used data from 17.0-public release and Biopharma Collaborative (BPC) v2.0-public for LUAD and GBM cohorts, respectively (downloaded 2025-06-02). The null prognostic model included baseline covariates, and the two constituent genomic features as main effects. The alternative model additionally included the PGI term. Baseline covariates included age and sex in all cohorts analyzed; tumor stage was additionally included for TCGA-LUAD and DFCI-LUAD, and CNA burden was additionally included for TCGA-LUAD and TCGA-GBM. PGIs with likelihood-ratio test (P < 0.05) were considered successfully validated. Because GENIE data was based on targeted panel sequencing, recurrent CNA features defined by GISTIC peak regions were not directly available. Therefore, we validated CNA-based genomic features using gene-level copy-number calls. A sample was called positive for a CNA feature if it harbored a concordant CNA (gain or loss, matching the TCGA-defined direction) in any gene located within the corresponding TCGA GISTIC peak region. Using this approach, we validated two PGIs: KRAS mutation with 9p21.3 loss in LUAD, and co-loss of 6p22.1 and 15q14 in IDH-wildtype GBM.

### Transcriptomic analysis of PGIs

We performed differential gene expression analyses for all PGIs using matched TCGA RNA-sequencing data. We applied DESeq2 with default settings using the TCGA gene-level count matrix as input. For each PGI within its corresponding TCGA cancer type, we modeled gene expression as a function of the PGI indicator while including the two constituent genomic feature indicators (G1 and G2) as covariates to account for their main effects. We extracted the log2 fold change and P-value for the PGI term to quantify the effect on gene expression associated with genomic co-alterations. P-values were adjusted for multiple testing using FDR. Established cancer genes highlighted in the analyses were collected from the Cancer Gene Census and the OncoKB Cancer Gene List.

### Pathway enrichment analysis of transcriptomic signatures of PGIs

To functionally interpret PGIs based on transcriptomic associations, we applied pathway enrichment analysis to the differential expression results for each PGI using the ActivePathways method (Paczkowska *et al*, 2020) with Holm family-wise error rate (FWER) correction for multiple testing. For each analysis, we removed lowly expressed genes from the background, defined as genes with median TPM < 1 in the corresponding TCGA cohort. We performed enrichment for all PGIs using the 50 MSigDB Hallmark gene sets (Liberzon *et al*, 2015). For the two clinically validated PGIs (KRAS mutation with 9p21.3 loss in LUAD; co-loss of 6p22.1 and 15q14 in IDH-wildtype GBM), we additionally performed enrichment using broader pathway collections.

Reactome pathways and Gene Ontology biological processes were retrieved from the g:Profiler software (Reimand *et al*, 2007) (downloaded 2026-03-24) and gene sets containing 50-250 genes were retained for the analysis, Benjamini-Hochberg FDR was used as the multiple testing correction method for a more permissive analysis, 0.05 as the cutoff parameter, and lowly expressed genes were removed from the background (TPM < 1), and default values for other parameters. Enrichment maps of significantly enriched pathways were generated in Cytoscape and were manually curated following standard protocols (Reimand *et al*, 2019).

### Clustering PGIs based on pathway enrichment profiles

We summarized each PGI by its pathway enrichment profile using the 50 hallmark gene sets. Each PGI was represented as a set of log-transformed adjusted P-values derived from the pathway enrichment analyses. We applied Uniform Manifold Approximation and Projection (UMAP) (McInnes *et al*, 2018) to the 50-dimensional vector of Hallmark enrichment values derived from PGI-associated differential expression results, to visualize PGIs and assess clustering based on pathway-level signatures. We then clustered PGIs in the UMAP embedding space using k-means with Euclidean distance. UMAP was performed in R using the umap package (v0.2.10.0) and clustering was performed using the NbClust R package (v3.0.1) (Charrad *et al*, 2014). To assess robustness, we explored a range of UMAP parameters, including the number of components (2-25) and the number of nearest neighbors (5-25), and repeated the analysis for all parameter combinations across 500 random seeds.

## Supplementary tables

**Table S1. Summary of cancer cohorts used for PGI discovery.** Overview of TCGA cohorts and sample sizes, detailing the specific features evaluated and the total PGIs identified within each cancer type.

**Table S2. Catalog of 57 PGIs.** Detailed information on PGIs, including constituent genomic features, sample counts, P-values and confidence intervals.

**Table S3. Enrichment analysis of MSigDB Hallmark gene sets.** Pathway enrichment analysis of 50 MSigDB Hallmark gene sets using the differential gene expression profiles of PGIs as input. Negative log10 of adjusted P-values are provided.

**Table S4. Dosage sensitivity and co-essentiality analyses of PGI-derived gene pairs.** The complete list of gene-gene pairs derived from each PGI based on dosage-sensitivity mapping in matching cancer transcriptomes is shown Further details about co-essentiality analysis of each pair are provided.

**Table S5. Differential expression analysis of the two selected PGIs in LUAD and GBM.** The output of differential expression analysis of the two PGIs in LUAD and GBM using DESeq2, including log-fold changes and adjusted P-values.

**Table S6. Integrative pathway enrichment analysis of the two selected PGIs in LUAD and GBM.** Results from the integrative pathway enrichment analysis using ActivePathways which utilized differential gene expression signatures from the two PGIs as input scores, and Reactome and Gene Ontology biological process as gene sets.

## Acknowledgments

This work was partially supported by the Canadian Institutes of Health Research (CIHR) Project Grant (PJT-197925) and Catalyst Grant (DV1-197665) to J.R., and the Investigator Award to J.R. from the Ontario Institute for Cancer Research (OICR). M.B. was partially supported by a Caven Fellowship, Excellence Award from the University of Toronto Department of Medical Biophysics, and a Doctoral Completion Award from the University of Toronto. L.L. was partially supported by mobility funding from the Kristjan Jaak National Scholarship Programme of Estonia. Funding to OICR is provided by the Government of Ontario. M.F. was supported by the Spanish State Research Agency (Agencia Estatal de Investigación, AEI) grant FPI PRE2021-101048. We would like to thank all patients and their families for donating tumor samples that enabled this research project. Results shown here are in whole or part based upon data generated by the TCGA Research Network: https://www.cancer.gov/tcga.

## Author Contributions

M.B. led data analyses and visualisation. J.R. conceptualized the project. M.B. and J.R. developed the methodology. J.R. and M.B. interpreted the data and wrote the manuscript. J.M., L.L. and M.F. contributed to data analyses and interpretation. J.R. supervised the project and acquired funding. All coauthors reviewed and approved the final draft of the manuscript.

## Supplementary materials

Masroor Bayati *et al*. (2026)

**Figure S1.**
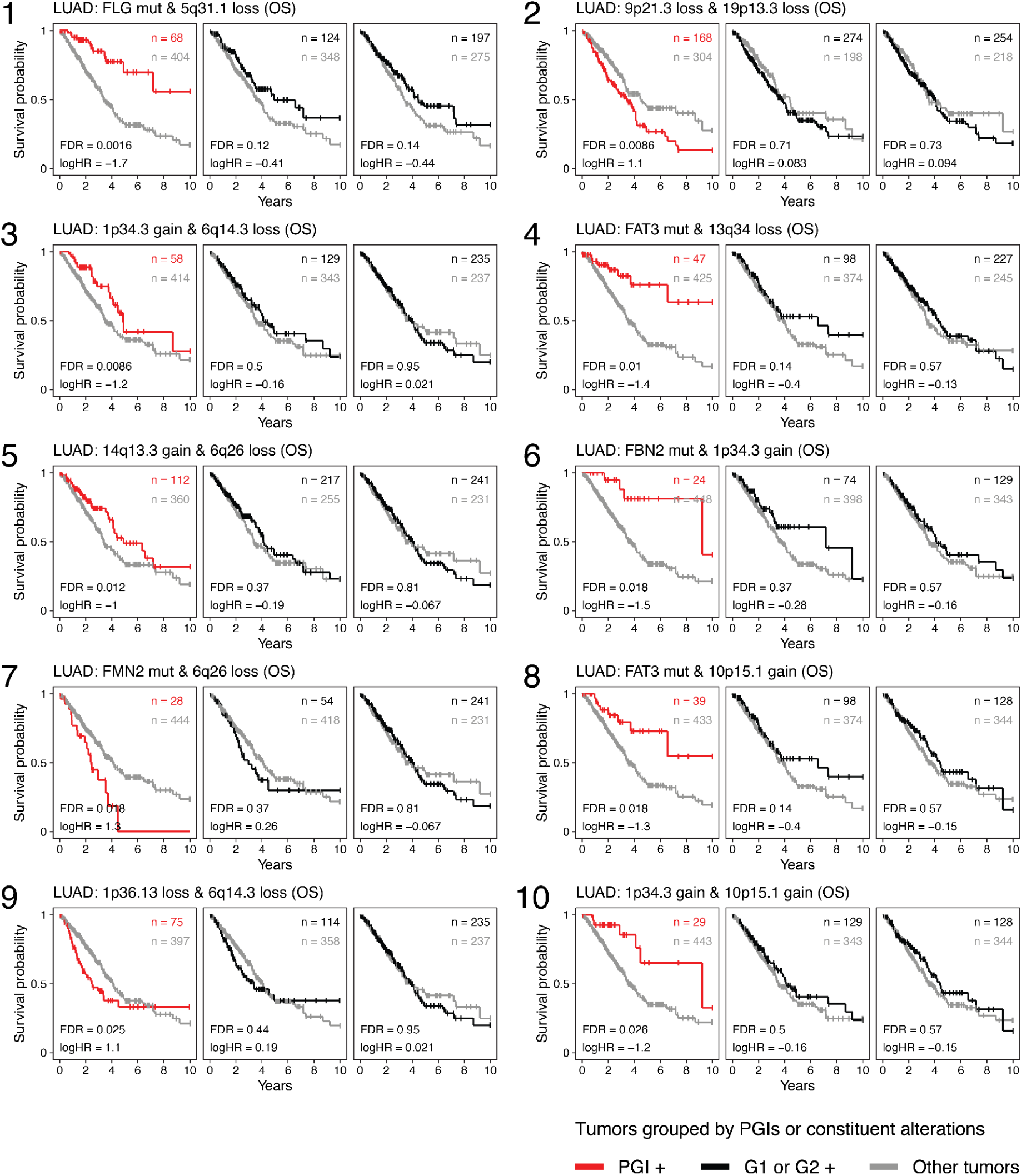
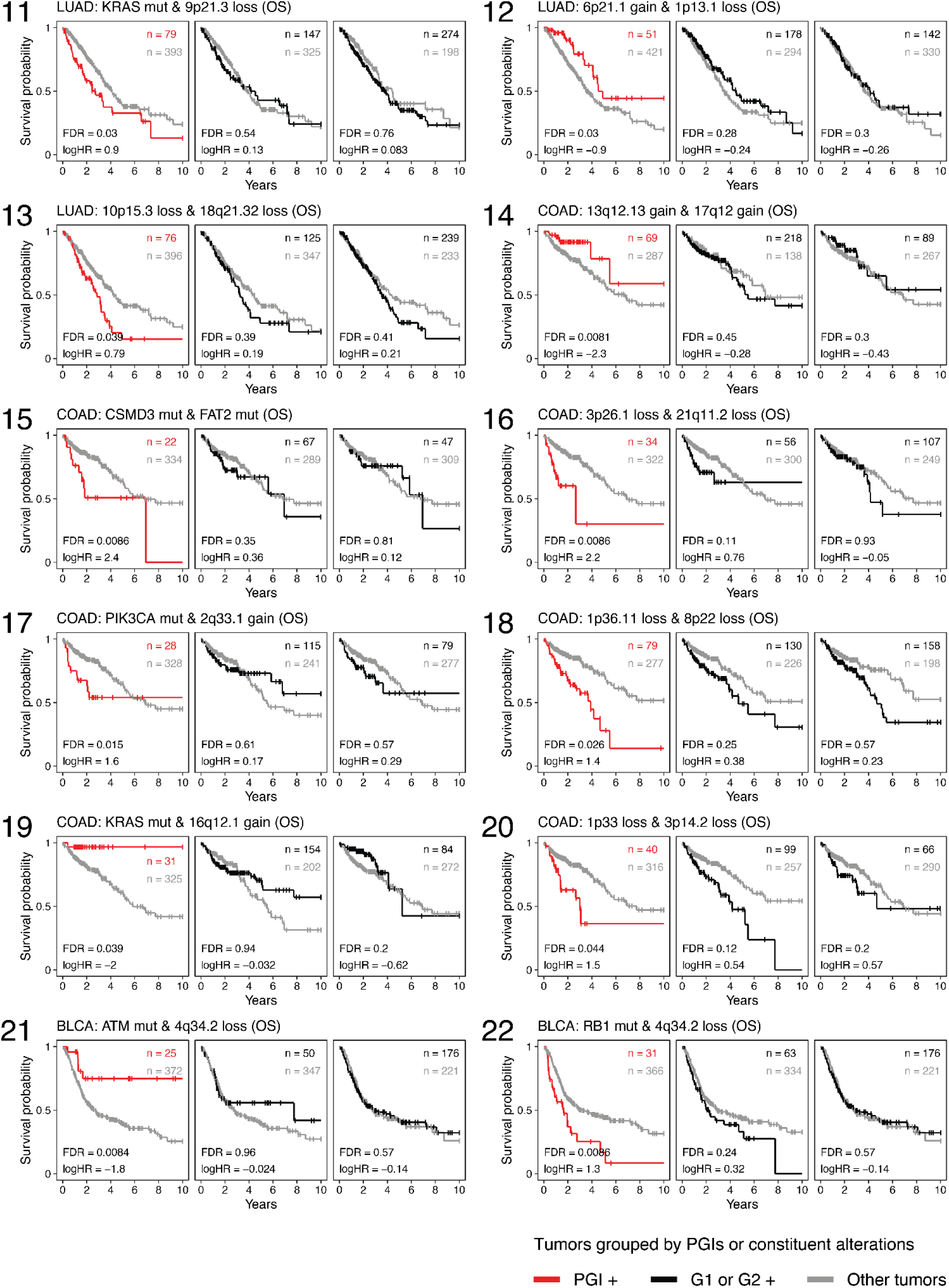
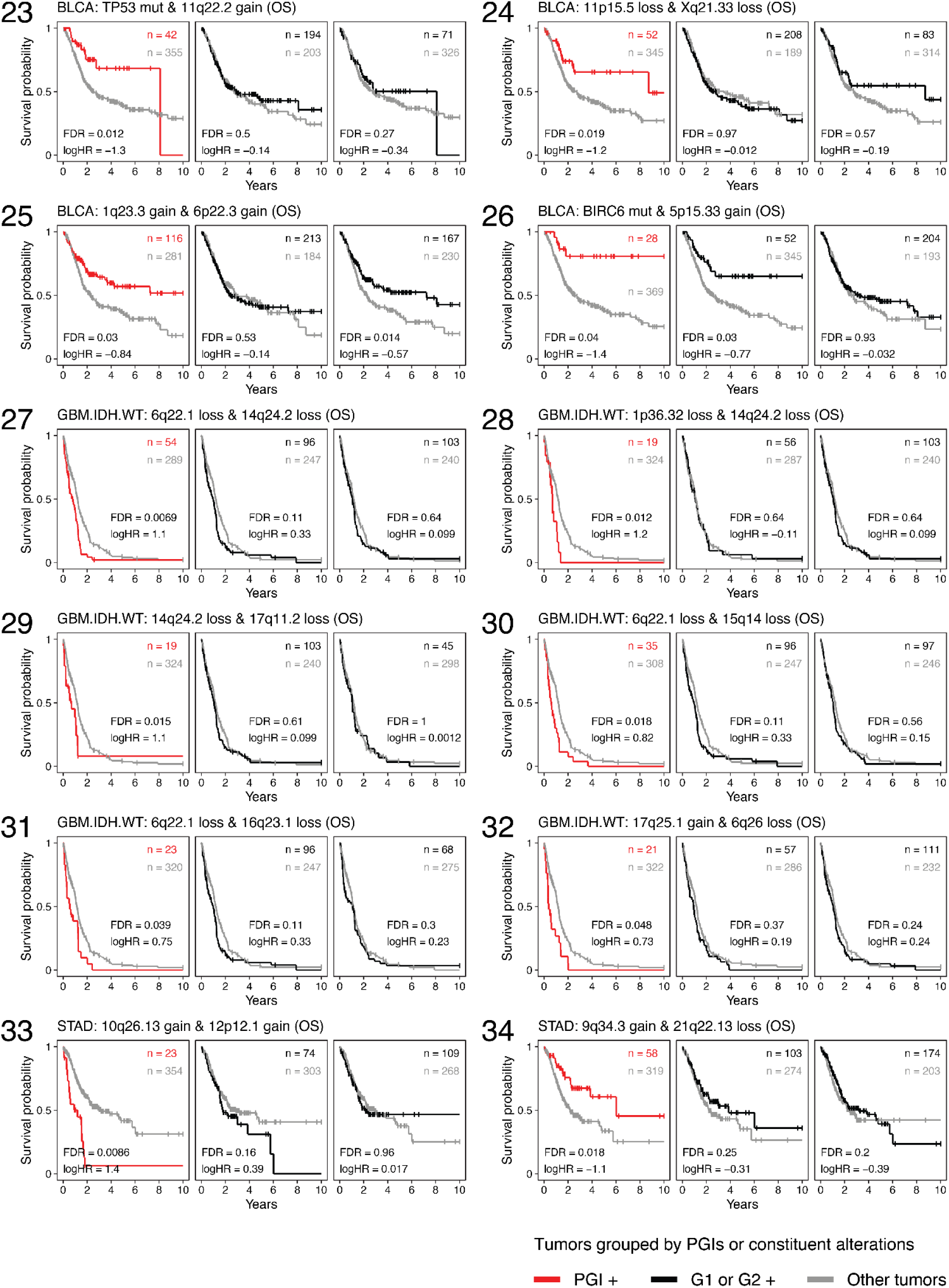
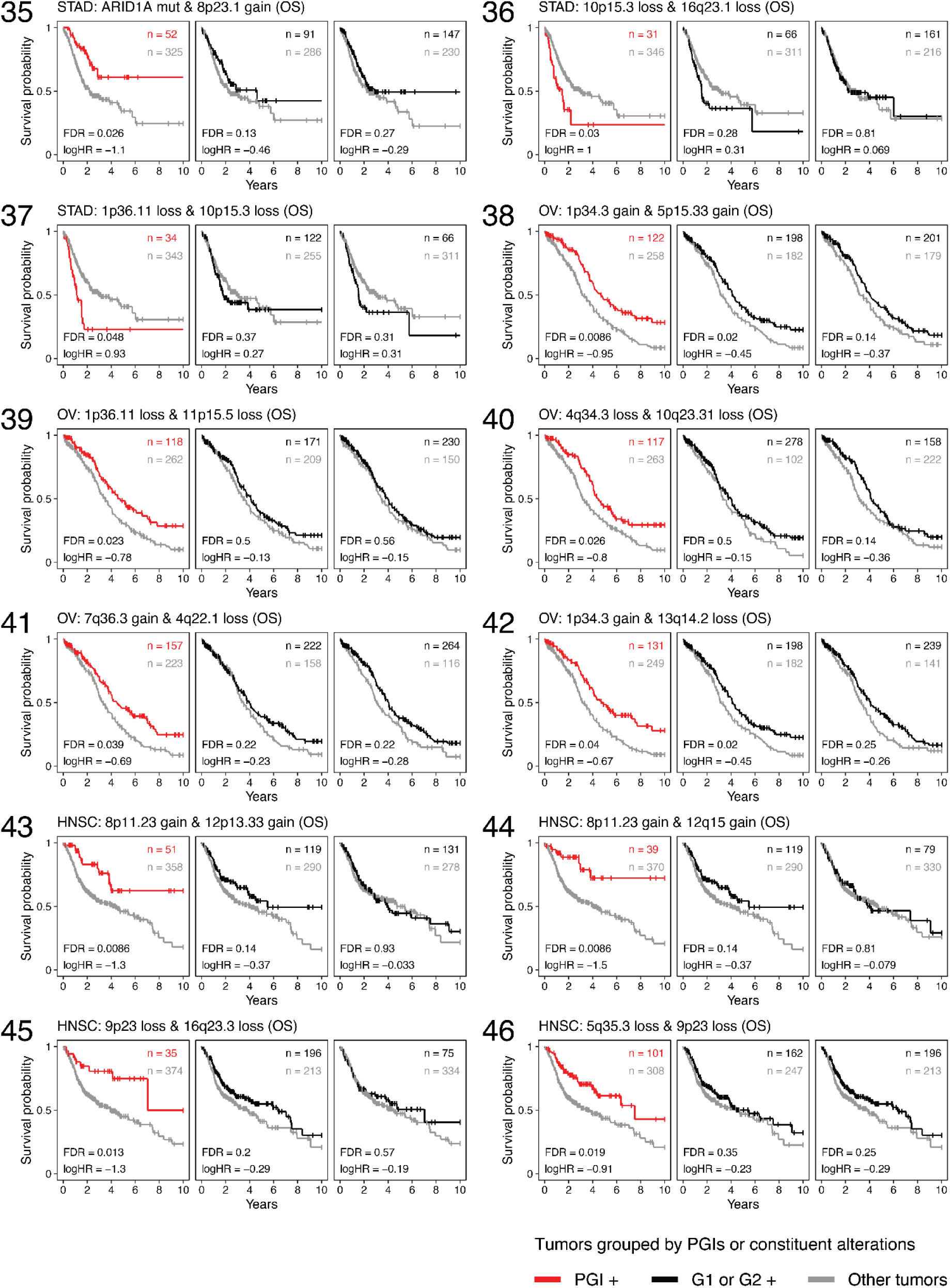
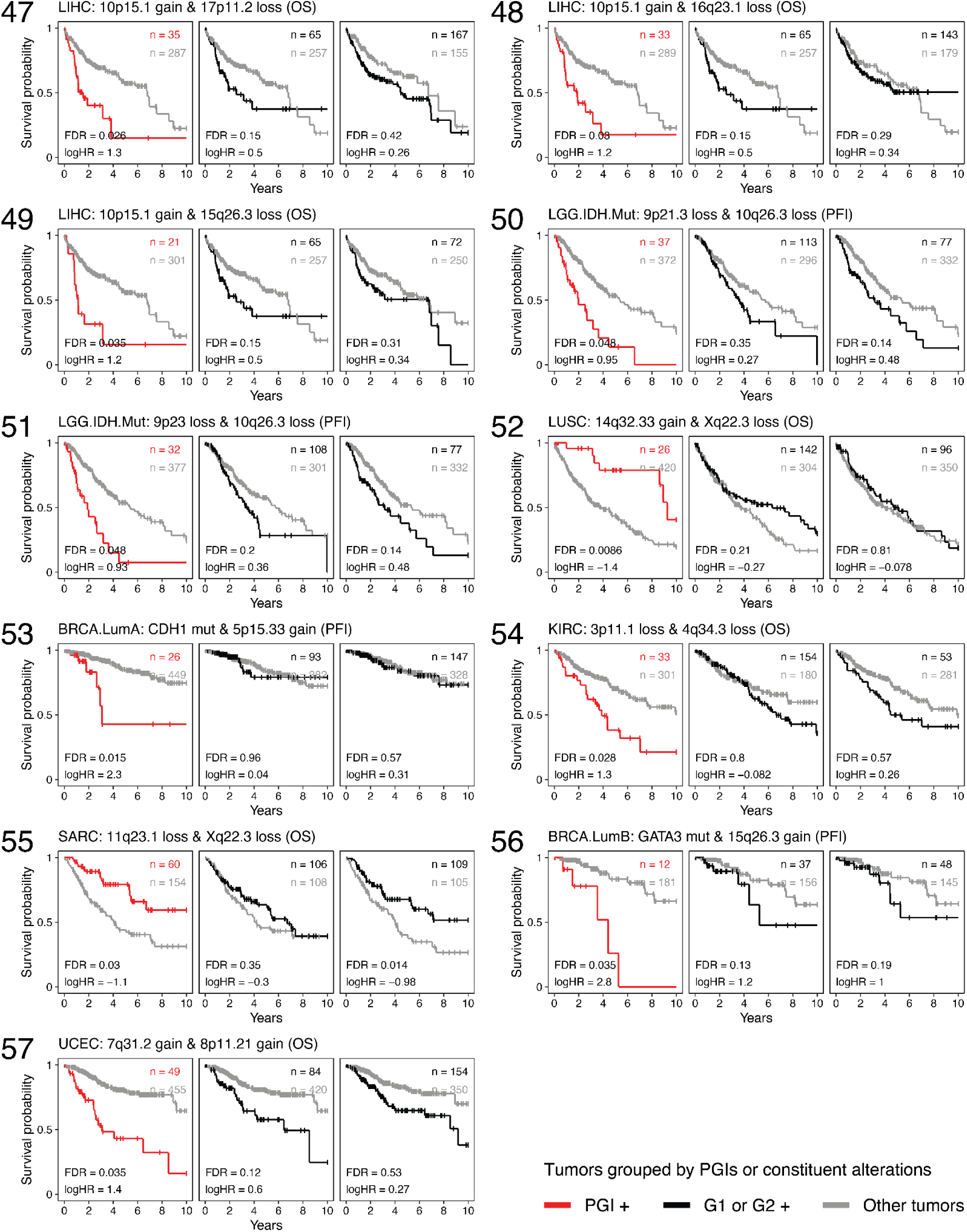
Catalog of PGIs with survival associations. Kaplan-Meier (KM) curves compare survival for PGI-positive versus PGI-negative patients (left), alongside survival stratified by each constituent genomic feature alone (G1 and G2) as controls (middle and right). Hazard ratios (HRs) and FDR-adjusted p-values were derived from CoxPH models adjusted for baseline clinical covariates, with PGI models further including G1 and G2 as covariates. Overall survival (OS) or progression-free intervals (PFI) are shown for different cancer types.

